# Precise electronic control of redox reactions inside *Escherichia coli* using a genetic module

**DOI:** 10.1101/2020.04.01.020511

**Authors:** Moshe Baruch, Sara Tejedor-Sanz, Lin Su, Caroline M. Ajo-Franklin

**Affiliations:** Department of BioSciences, Rice University, Houston, TX, 77005; Institute for Biosciences and Bioengineering, Rice University, Houston, TX, 77005

**Author notes:** Corresponding author: Dr. Caroline Ajo-Franklin, 6100 Main St., MS 140, Houston TX, 77005.

**Keywords:** microbial electrochemical technologies, electrobiosynthesis, microbial electrosynthesis, electrically-assisted fermentation

## Abstract

Microorganisms regulate the redox state of different biomolecules to precisely control biological processes. These processes can be modulated by electrochemically coupling intracellular biomolecules to an external electrode, but current approaches afford only limited control and specificity. Here we describe specific electrochemical control of the reduction of intracellular biomolecules in *Escherichia coli* through introduction of a heterologous electron transfer pathway. *E. coli* expressing *mtrCAB* from *Shewanella oneidensis* MR-1 consumed electrons directly from a cathode when fumarate or nitrate, both intracellular electron acceptors, were present. The fumarate-triggered current consumption occurred only when fumarate reductase was present, indicating all the electrons passed through this enzyme. Moreover, MtrCAB-expressing *E. coli* used current to stoichiometrically produce ammonia. Thus, our work introduces a modular genetic tool to reduce a specific intracellular redox molecule with an electrode, opening the possibility of electronically controlling biological processes such as biosynthesis and growth in any microorganism.

---

Microorganisms accomplish important biological functions such as conserving energy, regulating gene expression, and powering biosynthesis using different redox-active biomolecules. To enable control of these processes in any microorganism, researchers have coupled the redox state of these biomolecules to an external electrode using membrane-permeable, small molecule redox mediators^1–5^, redox polymers^6^, and membrane-intercalated nanostructures ^7,8^. These approaches can allow cells to produce electrical current or consume it, resulting in either oxidation or reduction of intracellular redox species, respectively. Bioelectrochemical devices can then be used to drive biosynthetic reactions ^1,4,5^, perform bioelectronic sensing ^9^, actuate gene expression^3^, and modulate cellular growth ^10,11^ within the microorganism of interest. Despite these accomplishments, these strategies couple the redox state of the electrode to multiple intracellular redox biomolecules, resulting in off-target effects, cellular toxicity, and poor control of biosynthesis ^1,3,4^. To achieve precise electrochemical control of a biological process, a strategy that couples an electrode to a specific intracellular redox pool is still needed ^12,13^.

To couple an electrode to specific redox molecules in a bacteria of our choosing, we and others have introduced genes from the Mtr pathway from *Shewanella oneidensis* MR-1 into heterologous bacterial hosts^14–18^. Under anaerobic conditions, *S. oneidensis* can use the Mtr pathway to transfer electrons from catabolism to produce a current at an extracellular electrode (**Figure 1A**). Electrons released from catabolism of lactate are first transferred from NADH to menaquinone (MK) by Complex I ^19^. From MK, electrons traverse the cell envelope through a series of multiheme cyts *c* ^20^: CymA in the inner membrane ^21^, Stc in the periplasm^22^, across the outer membrane via the MtrCAB complex^23^, and directly from MtrC to an anode ^24^, typically biased at +200 mV_Ag/AgCl_. Interestingly, the Mtr complex also permits current consumption from a cathode biased at +560 mV_Ag/AgCl_ to intracellular fumarate under aerobic conditions ^25,10^. In this case, electrons are transported to MtrC, across the outer membrane by the MtrCAB complex, to CymA, to the MK, and to the periplasmic fumarate reductase, FccA ^25^ ultimately reducing fumarate (**Figure 1B**). This current consumption opens the possibility to use electricity to reduce CO_2_ and N_2_ to fuels and ammonia, respectively, and an area of very active research^26,27^.

**Figure 1.**
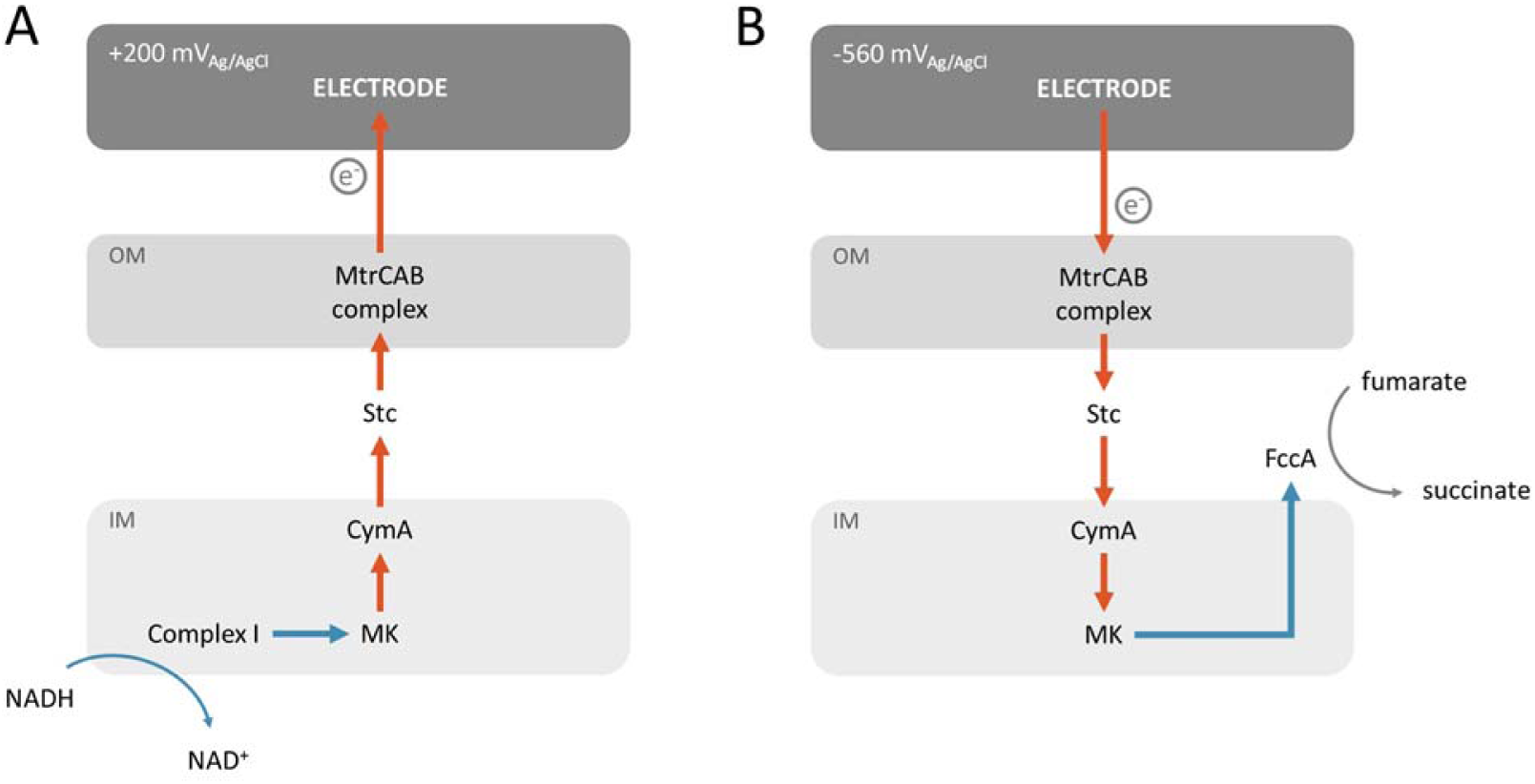
Coupling of intracellular redox reactions to an electrode in *Shewanella oneidensis* MR-1. Schematic illustrating the role of the MtrCAB complex and the inner membrane cyt *c* CymA and menaquinone (MK) in the coupling of current production to intracellular oxidation of NADH (A) and current consumption (B) to intracellular reduction of fumarate in *S*.*oneidensis* MR-1. (OM: outer membrane, IM: Inner membrane).

We have previously shown that oxidation of intracellular lactate can be coupled to current production in *Escherichia coli* that heterologously express *cymAmtrCAB* ^16,18^. This led us to hypothesize that the Mtr pathway could couple oxidation of a cathode to reduction of intracellular biomolecules. Here we probe whether Mtr-expressing *E. coli* can directly consume current from a cathode, compare the route of electron flow under anodic and cathodic conditions, and show for the first time that an electrode can stoichiometrically drive the reduction of a specific molecule inside engineered *E. coli*.

## Results

### *E. coli* consume current using *mtr* and native oxidoreductases

We first sought to determine if the Mtr pathway could allow cathodic electrons to directly enter a heterologous host upon addition of an electron acceptor. Since *E. coli* has two MK-linked fumarate reductases, FrdABCD and SdhABCD, we hypothesized that the Mtr pathway in *E. coli* could deliver cathodic electrons via these native proteins to fumarate (**Figure 1A**). To probe the specific role of the Mtr pathway, we compared the behavior of several strains: *E. coli* expressing only the cytochrome *c* maturation (*ccm*) genes (abbrev. Ccm-*E*.*coli*)^17^, *E. coli* expressing *ccm* and *mtrCAB* (abbrev. Mtr-*E. coli)*^15^, and *E. coli* expressing *ccm* and *cymAmtrCAB* (abbrev. CymAMtr-*E. coli)*^17^. The *ccm* genes are required to make cytochromes *c* in the C43(DE3) parental background.

To prepare *E. coli* for cathodic conditions, individual strains were first grown aerobically, incubated in potentiostatic-controlled bioreactors under anaerobic, anodic conditions (ΔV=+200 mV_Ag/AgCl_) for at least one day. Under these conditions, the CymAMtr*-E. coli* strain produced a significant steady-state current, while Ccm*-E. coli* and Mtr*-E. coli* produced much lower currents (**Figure 2A**), reinforcing that CymA is important for current production^16–18^. As anaerobic conditions were maintained, the electrode bias was then switched to cathodic conditions (ΔV= −560 V_Ag/AgCl_), fumarate was added, and current consumption was measured. In the absence of *E. coli*, neither current consumption nor fumarate reduction was observed (**Figure S1A**). Likewise, the Ccm-*E. coli* strain did not consume significant levels of current (**Figure 2B**). In contrast, both the Mtr-*E. coli* and the CymAMtr-*E. coli* strains consumed significant levels of current (**Figure 2B**), starting within 30 seconds after fumarate addition (**Figure S1B**). This rapid onset indicates that a change in gene expression was not needed for current consumption. There is no significant difference in the MtrCAB abundance in the CymAMtr-*E. coli* and Mtr-*E*.*coli* ^17^, so we can rule out a difference in gene expression as the origin for the current difference. Thus, while CymA is required for current consumption in *S. oneidensis*, it is not required for current consumption in *E. coli*. NapC, which can complement CymA, is disrupted in the C43(DE3) background ^28^, so it cannot be involved in current consumption. Rather, MtrCAB either directly or more probably, indirectly through as-yet-unknown native biomolecule inside *E. coli*, enables new host microorganisms to directly accept electrons from a cathode. To maintain optimal metabolic activity in the strains during the anodic acclimation, we used CymAMtr-*E. coli* in the rest of our experiments.

**Figure 2.**
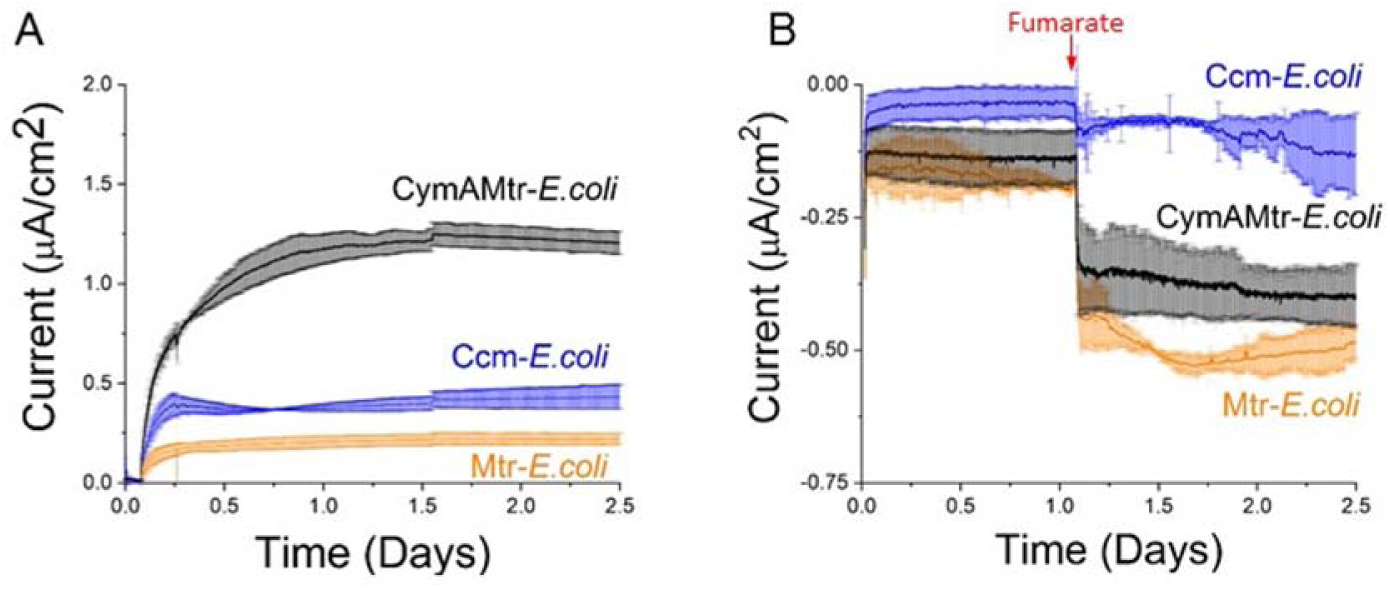
Expression of *mtrCAB* from *S. oneidensis* MR-1 allows *Escherichia coli* to directly produce or consume current. (A) Chronoamperometry of bioelectrochemical reactors containing Ccm-*E. coli* (orange), Mtr-*E. coli* (orange), or CymAMtr-*E. coli* (black) in the anodic compartment with the anode poised to +200 mV_Ag/AgCl_. Lactate is provided as an electron donor and the anodic chamber is kept anaerobic by bubbling with N_2_(g). (B) Chronoamperometry of bioelectrochemical reactors containing Ccm-*E. coli* (orange), Mtr-*E. coli* (orange), or CymAMtr-*E. coli* (black) in the cathodic compartment with the cathode poised to −0.56V_Ag/AgCl_. Addition of fumarate is indicated by the red arrow. The bars indicate the standard deviation in current from three bioreactors.

Cyclic voltamettry of the CymAMtr-*E. coli* strain with fumarate (**Figure S1C**) revealed negative shift of the catalytic wave starts at −43 mV vs. Ag/AgCl, which is close to the redox potential of FrdAB^29^, suggesting that electrons entering the Mtr pathway could transfer to the *E. coli* fumarate reductase. To determine whether all cathodic electrons passed through the native fumarate reductases of *E. coli* upon fumarate addition, we compared the amount of current consumed by CymAMtr-*E. coli* in the wt, *ΔfrdABCD* (abbrev. Mtr-*Δfrd*), and *ΔfrdABCDΔsdhABCD* (abbrev. Mtr-*ΔfrdΔsdh*) backgrounds. The CymAMtr-*Δfrd* strain consumed ∼40% as much current as the CymAMtr-*E. coli* strain (**Figure 3A**) and the CymAMtr-*ΔfrdΔsdh* strain did not consume any significant current (**Figure 3B**). These observations strongly suggest that cathodic-derived electrons pass solely through FrdABCD and SdhABCD upon fumarate addition in *E. coli* expressing *mtrCAB*.

**Figure 3.**
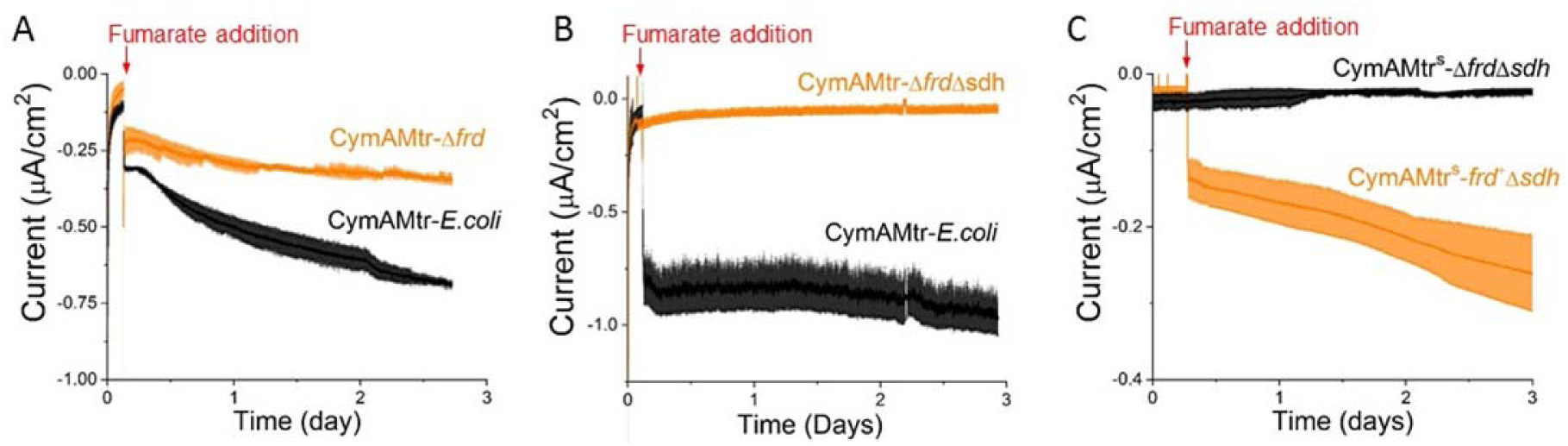
Fumarate-triggered current consumption in CymAMtr-*E. coli* requires *E. coli* fumarate reductases. (A) Current consumption by the CymAMtr-*E. coli* in the wt (black) and *ΔfrdABCD* (orange) background upon addition of fumarate, showing reduced current consumption in the *ΔfrdABCD* background. (B) Current consumption by CymAMtr-*E. coli* in the wt (black) and *ΔfrdABCD ΔsdhABCD* (orange) backgrounds upon addition of fumarate, showing that current is not consumed when fumarate reductase is absent. (C) Current consumption by CymAMtr-E. coli in the Δ*frdABCD* Δ*sdhABCD* (black) and *frdABCD-*complemented Δ*sdhABCD* background (orange). All the experiments were performed under anaerobic conditions with the cathode poised to −560 mV_Ag/AgCl_. Addition of 50mM fumarate is indicated by the red arrow, and the bars indicate the standard deviation in current from three bioelectrochemical reactors.

To confirm that the inability of CymAMtr-*ΔfrdΔsdh* to uptake electrons was not a polar effect of the *frd* and *sdh* deletions, we probed current consumption in strains with complemented expression of *frdABCD*. To do so, we first altered the regulation of the *cymAmtrCAB* operon to accommodate expression of *frdABCD* from a third plasmid (see **Supplementary Information** for more details). This created the parental CymAMtr^S^-*ΔfrdΔsdh* strain and the complemented CymAMtr^S^-*frd*^*+*^*Δsdh* strain, which could reduce fumarate and express CymA MtrCAB (**Figure S2C**). The CymAMtr^S^-*frd*^*+*^*Δsdh* strain consumed a significant current upon fumarate addition (**Figure 3C**), in contrast to the CymAMtr^S^-*ΔfrdΔsdh* strain, which did not consume any current. These data rule out the possibility that polar effects in the Mtr^S^-*frd*^*+*^*Δsdh* background are responsible for reduced current consumption and show the Mtr pathway delivers cathodic electrons via the MK-linked fumarate reductases to fumarate in *E. coli*. More broadly, cathodic current flows through only FrdABCD in the Mtr^S^-*frd*^*+*^*Δsdh* upon addition of fumarate, which is the first demonstration to our knowledge of a heterologous genetic module that directs electrons to only a single native oxidoreductase inside a bacterial strain.

### Menaquinone and Complex I are essential for coupling intracellular oxidations to an anode, but not for coupling reductions to a cathode

Since a MK-linked fumarate reductase is essential for current consumption under cathodic conditions, it is likely that cathodic electrons flow through a quinone. To test this hypothesis, we examined the bioelectrochemical behavior of two strains expressing *cymAmtr* that lack genes essential for menaquinone synthesis, *menA*^30^ (abbrev. CymAMtr-Δ*menA)* and *menC* ^31^. *menC* is also essential for synthesis of the quinone-derived redox shuttle ACNQ ^32^, allowing us to also test whether ACNQ is an electron carrier here. As before, these strains were acclimated in bioelectrochemical reactors under anodic conditions with similar cell densities before being switched to cathodic conditions.

Under anodic conditions, current production by the CymAMtr-Δ*menC* and CymAMtr-Δ*menA* strains significantly declines to near the current levels produced by the Ccm-*E. coli* strain (**Figure 4A, C**). Complementation of the *menC* in trans restores the current production to the wt levels (**Figure 4C**). These observations demonstrate that menaquinone mediates electron flow from the cytosol to CymA in *E. coli* just as in *S. oneidensis* MR-1^33,34^. Under cathodic conditions, the current consumption by CymAMtr-expressing *E. coli* in the Δ*menA* and Δ*menC* backgrounds was not significantly different from the wt background (**Figure 4B, D**). These surprising observations indicate that the current consumption in Mtr-expressing *E. coli* does not rely on the presence of menaquinone or ACNQ, in contrast to *S. oneidensis* ^25,32^ and other reports in *E. coli* ^35^, and that the fumarate reductase of *E. coli* accepts electrons either directly from MtrA or indirectly through as-yet-unknown native biomolecule inside *E. coli*.

**Figure 4.**
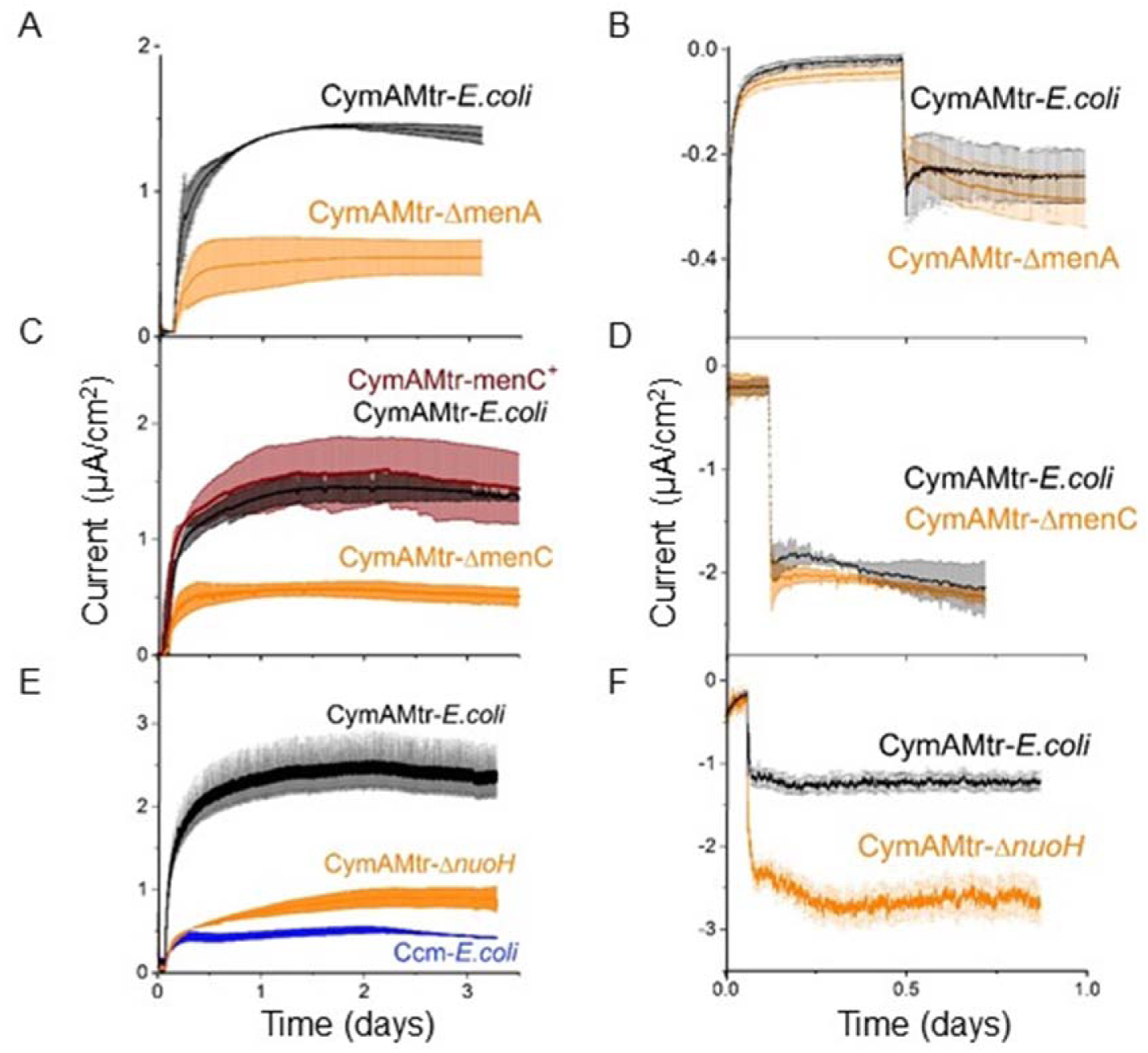
Mtr-expressing *E. coli* requires menaquinone and Complex I to generate current, but does not require them to consume current. A,C,E) Current production under anodic conditions and B,D,F) current consumption under cathodic conditions for the CymAMtr-*E. coli* in wt (black) and gene deletion (orange) backgrounds. A,B) Current as a function of time for CymAMtr-*E*.*coli* in the A,B) Δ*menA* background and C,D) *ΔmenC* background (orange). Complemented strain menC+ is shown in red. E) Current as a function of time for CymAMtr-*E*.*coli* in the Δ*nuoH* background. The anode was poised to +200 mV_Ag/AgC_ and the cathode was poised to −560 mV_Ag/AgCl_ for all experiments. Fumarate addition (50mM) is indicated by the red arrow, and the bars indicate the standard deviation in current from triplicate bioelectrochemical reactors.

While fumarate reductase accepts electrons from the Mtr pathway, we also assume it is still able to accept electrons from MKH_2_. Thus it is likely that these two pathways may compete for fumarate reductase, and eliminating MK reduction by catabolic reactions would increase current consumption. Lending credence to this hypothesis, we observed that the longer CymAMtr-*E. coli* was deprived of a carbon source in our bioreactors (and thus the lower the catabolic rate), the higher current consumption was upon fumarate addition (**Figure S3A**). Since Complex I (NDH-1) catalyzes the transfer of electrons from NADH to MKH_2_ under anaerobic conditions ^36^, we probed the effect of disrupting Complex I on current consumption. We prepared CymAMtr-*E. coli* lacking a functional NDH-1 ^37,38^, abbrev. CymAMtr-*ΔnuoH*, and acclimated this strain in bioreactors as before.

Under anodic conditions, the CymAMtr-*ΔnuoH* strain produced only ∼33% as much current as the CymAMtr-*E. coli* and only slightly more current than the Ccm-*E. coli* (**Figure 4E**) Under cathodic conditions, the CymAMtr-*ΔnuoH* strain consumed ∼225% more current than the CymAMtr-*E. coli* upon fumarate addition (**Figure 4F**). Heme staining of whole cell lysates of CymAMtr-*ΔnuoH* showed that expression of the Mtr cyt *c* was significantly lower in the deletion strain (**Figure S3B**) compared to the CymAMtr-*E. coli*. This observation rules out that higher levels of Mtr cyt *c* in the *ΔnuoH* background causes the increased current consumption. Instead, these data strongly suggest that eliminating a competing electron flux into fumarate reductase allows additional cathodic electrons to enter the Mtr pathway.

### Current consumption by Mtr-expressing E. coli yields non-stoichiometric accumulation of succinate

Having demonstrated that current consumption in *E. coli* requires *mtrCAB* and fumarate reductase, we turned to the question of whether cathodic electrons transported by the Mtr pathway could stoichiometrically drive intracellular reduction of fumarate to succinate. We monitored the CymAMtr-*ΔnuoH* strain in reactors poised to cathodic conditions (polarized reactors) and into reactors which were not connected (unpolarized reactors) and measured the extracellular concentrations of several organic acids after addition of fumarate. We found it necessary to supplement the reactors with 40 mM pyruvate, a fermentable carbon source, to sustain bacterial viability during the 14-day long experiment. Without fumarate, pyruvate by itself did not trigger any current consumption (Data not shown) and did not introduce additional electrode-coupled reactions in bioelectrochemical reactors (**Figure S5A**).

The polarized reactors steadily consumed current and accumulated succinate at a different rate than the unpolarized reactions (**Figure 5A**). Knowing that cathodic electrons only reduce fumarate via fumarate reductases (**Figure 2**) and assuming that succinate consumption is equal under polarized and unpolarized conditions, we used the average current consumption to estimate that an additional 0.49 mM succinate would accumulate in the polarized reactors over 14 days compared to unpolarized reactors. However, we observed that the polarized reactors accumulated 29% less succinate than the unpolarized reactors, 12.63 ±1.68 mM vs 20.28 ±0.97 mM over 14 days, respectively. Thus, the number of electrons accumulated in succinate is opposite in direction and 10 fold-higher in magnitude than what we expected. Overall, the concentrations of other organic acids we monitored were very similar in the polarized and unpolarized reactors (**Figure S5C**). Only the formate concentration was slightly higher in the polarized reactor by 2.38 ±0.05 mM after 9 days (**Figure S5C**), but this minor change is insufficient to explain the dramatic difference between the expected and observed succinate accumulation. Surprisingly, when we repeated this experiment with the CymAMtr-*E. coli* strain we did not detect any significant difference between the polarized and unpolarized reactors (**Figure S5D**) suggesting that the higher current consumption by the CymAMtr-*ΔnuoH* is essential for this phenotype.

**Figure 5.**
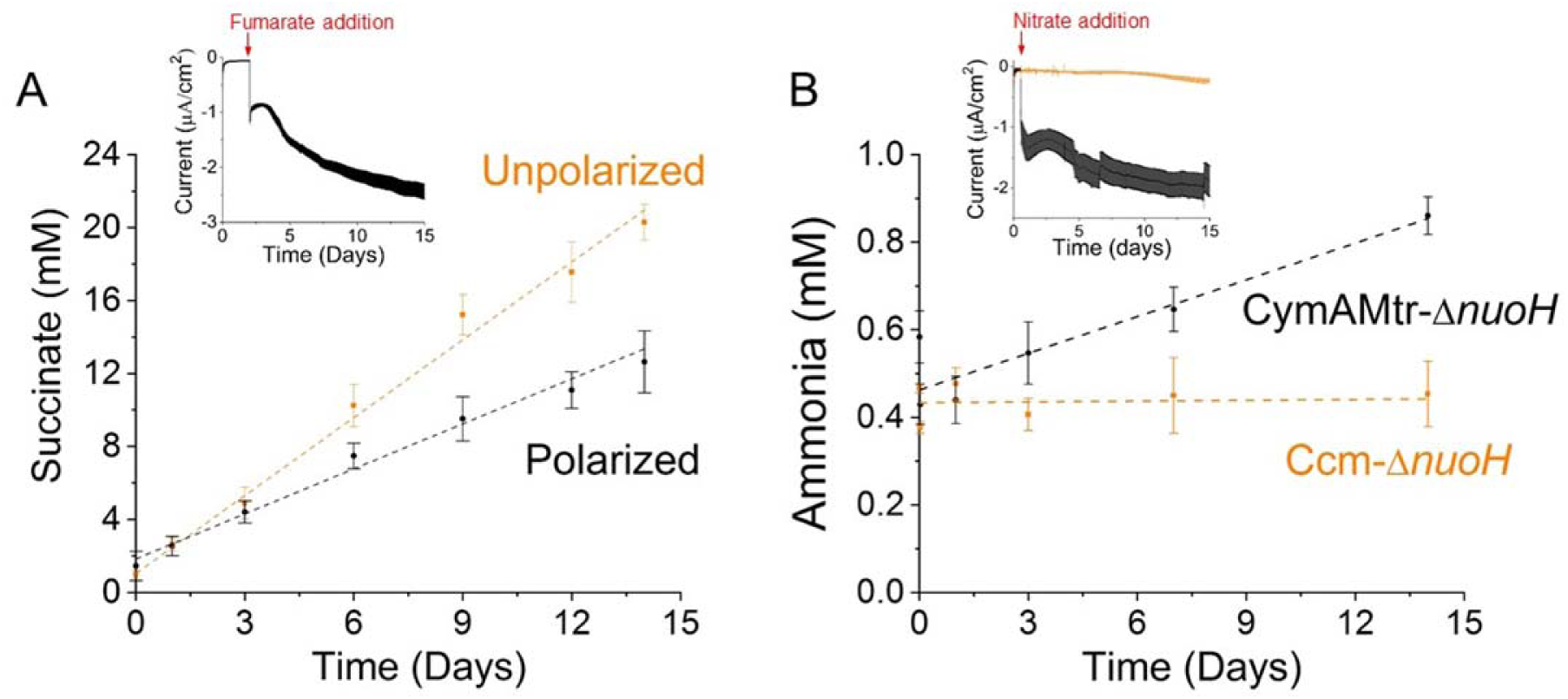
Current consumption is coupled to non-stoichiometric accumulation of succinate, but stoichiometric accumulation of ammonia. A) The concentration of succinate in the extracellular media as a function of time in polarized (black) and unpolarized (orange) bioelectrochemical reactors containing the CymAMtr-*ΔnuoH* strain. Dashed lines indicate the linear trends for the polarized (black, R^2^ −0.98) and unpolarized (orange, R^2^ −0.99) reactors, displaying a slope of 0.82 mM succinate day^-1^ and 1.42 mM succinate day^-1^, respectively. (inset) Current consumption in the polarized reactors upon fumarate addition. B) The concentration of ammonia in the extracellular media as a function of time in bioelectrochemical reactors containing the CymAMtr-*ΔnuoH* strain (black) or Ccm-*ΔnuoH* (orange) after addition of 10 mM nitrate. Dashed lines indicate the linear trend for the ammonia production by CymAMtr-*ΔnuoH* (black, R^2^ −0.88) displaying a slope of 0.027 mM ammonia day^-1^. The ammonia concentration for the Ccm-*ΔnuoH* strain (orange line provided to guide the eye) does not change significantly. In all experiments, the cathode is poised to −560 mV_Ag/AgCl_ and the bars indicate the standard deviation in current from triplicate bioelectrochemical reactors.

These observations, layered upon our knowledge that fumarate is only being reduced by fumarate reductases in Mtr-expressing *E. coli*, led us to question the assumption that succinate was equal in the polarized and unpolarized bioelectrochemical reactors. Fumarate and succinate are both intermediates in the TCA cycle and are acted on by enzymes that are both allosterically and transcriptionally regulated. Moreover, transcriptional regulation is governed by the redox state of the cell, making it very possible that there is differential regulation in the polarized and unpolarized conditions.

### Current consumption by Mtr-expressing E. coli yields stoichiometric accumulation of ammonia

Our postulate that non-stoichiometric accumulation of succinate was due to unequal consumption led us to examine whether intracellular reductions could be stoichiometrically driven via other oxidoreductases where the product is not utilized under our conditions. Since we expect ammonia will not be utilized under our conditions, and the Nar and Nir enzyme complexes that catalyze the reduction of nitrate ammonia are located in the inner membrane ^39,40^, we chose to examine reduction of nitrate to ammonia. In the C43(DE3) background, this reduction is a two-step reaction: the two-electron reduction of nitrate to nitrite by Nar or Nrf is followed by the six-electron reduction of nitrite to ammonia by Nir. The *nap* operon in C43(DE3) is non-functional ^28^.

To establish whether nitrate and nitrite could be reduced by the Mtr pathway in *E. coli*, we added nitrate or nitrite to reactors without bacteria, with Ccm-Δ*nuoH*, and with CymAMtr-*ΔnuoH* and monitored current flow. Nitrite is toxic to *E. coli* at high concentrations, so small aliquots of nitrite were added periodically in these experiments to keep its steady-state concentration low. No current was consumed in the reactors without bacteria, confirming that these molecules were not abiotically reduced. While low levels of current were consumed upon either nitrate (**Figure 5A**) or nitrite (**Figure 5B**) addition to reactors containing the Ccm-*E. coli*, the CymAMtr-*E. coli* consumed ∼5.6 fold and ∼2.5 fold higher current levels, respectively, indicating that the majority of the cathodic electron flux in CymAMtr-*E. coli* is Mtr-dependent. These data provide an additional example of delivery of electrons to inner membrane oxidoreductases in a heterologous host by *mtrCAB*.

To eliminate the effect of the small amount of current that is consumed independently of the Mtr pathway, we compared the accumulation of ammonia in bioreactors containing the Mtr-*ΔnuoH* strain and the Ccm-*ΔnuoH* strain after nitrate addition. Based on the difference in current consumption between the Mtr-*ΔnuoH* and the Ccm-*ΔnuoH* strains (**Figure 5A)**, we predicted that stoichiometric utilization of cathodic current would cause the Mtr-*ΔnuoH* strain to accumulate between 0.123 mM to 0.493 mM more ammonia over 14 days compared to the Ccm-*ΔnuoH strain*. (The lower bound assumes that cathode current reduces nitrate completely to ammonia, while the upper bound assumes that cathodic current contributes only to nitrite production. See SI for additional details.) Indeed, the Mtr expressing strain started accumulating more ammonia immediately at a significantly faster rate than the negative control (**Figure 5C**), and over 14 days, accumulates 0.405 mM more ammonia (**Figure 5C**). The excellent agreement between the observed and expected changes in ammonia accumulation indicate that cathodic electrons delivered through the Mtr pathway were used to stoichiometrically reduce nitrate to ammonia. More broadly, these data indicate that *mtrCAB* is a genetic module that can be used to drive specific, highly reductive biotransformations within novel microorganism hosts.

## Discussion

Here we show that the Mtr pathway can specifically deliver electrons to intracellular oxidoreductases and can drive intracellular redox reactions in a stoichiometric manner. Upon fumarate addition, *E. coli* take up electrons from the cathode via the MtrCAB complex and pass them to the fumarate reductases (**Figure 6**). The amount of current consumed can be increased by eliminating the MK reductase, Complex I (**Figure 4**). Interestingly, the current consumed is not stoichiometrically related to the accumulation of succinate from succinate, but is stoichiometric with reduction of nitrate to ammonia (**Figure 5**). Taken together, this work demonstrates use of *mtrCAB* as a genetic module to stoichiometrically drive specific intracellular redox reactions in heterologous hosts. Below, we discuss the implications of this work for designing genetic modules for coupling cathode oxidation to intracellular reductions and opportunities for modulating cell behavior.

**Figure 6.**
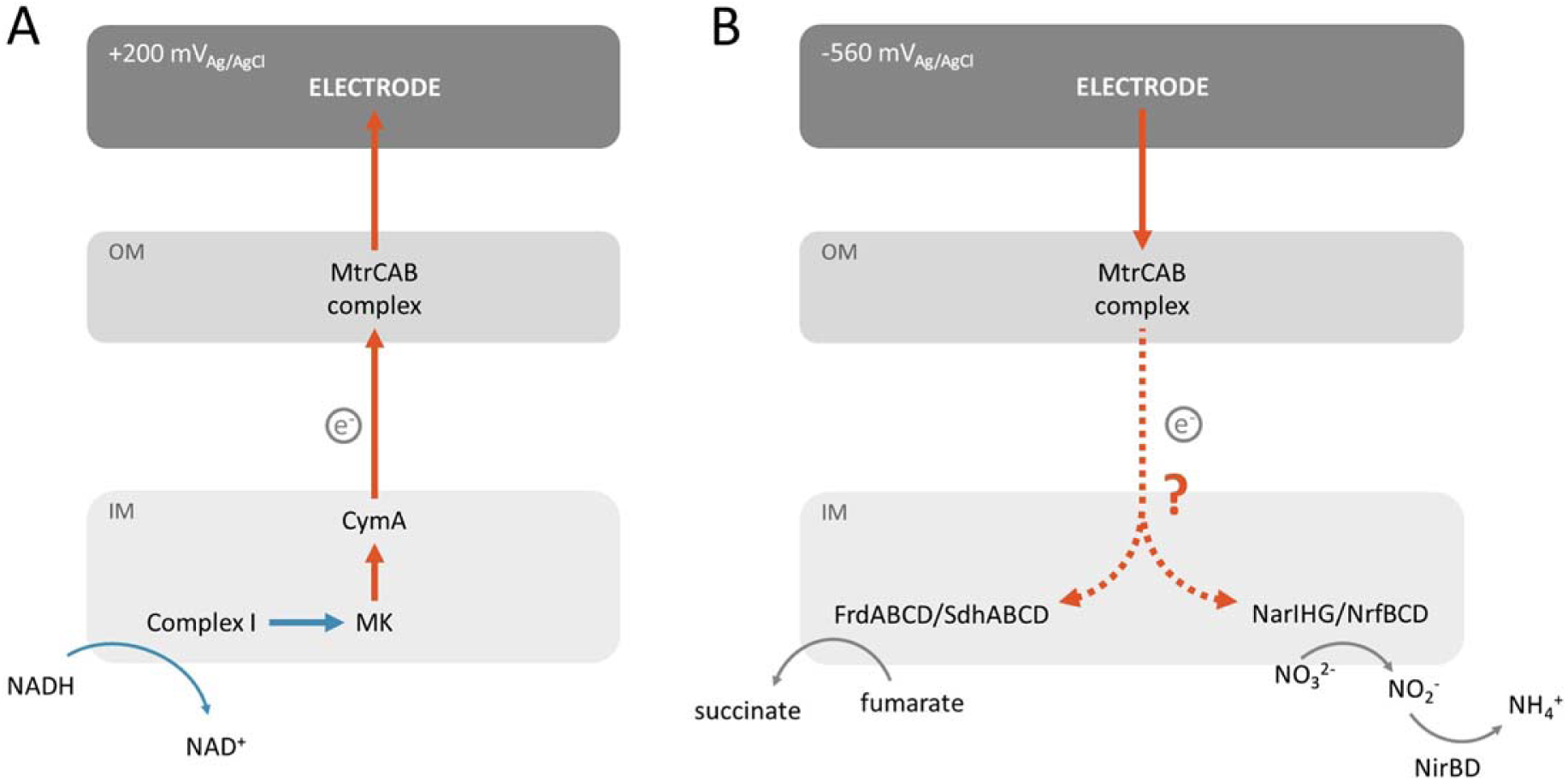
Model for how heterologously expressed MtrCAB couples intracellular oxidations (left) and reductions (blue arrows) to current production and consumption, respectively, in *E. coli*. (A) Oxidation of NADH by Complex I transfers electrons to MK. Electrons flow from MK via CymA to the MtrCAB complex. (B) MtrCAB transfer electrons via an unknown process to FrdABCD/SdhABCD and the NarIHG/NrfBCD enzymes, which in turn reduce fumarate and nitrate to succinate and nitrite, respectively. Further reduction for the nitrite to ammonia is mediated through NirBD.

### The Mtr pathway is a genetic module to specifically alter cell behavior by driving intracellular reactions stoichiometrically

The finding that the Mtr pathway can be used to drive reductions by inner membrane oxidoreductases provides new opportunities to power key biological processes with electricity in a variety of microorganisms. For example, since the chemolithoautotroph *Nitrosomonas europaea* can use ammonia as its sole energy source and reductant ^41^, the production of ammonia via electricity, Mtr, and nitrate/nitrite reductases could be used to produce *N. europaea* biomass. Alternatively, we envision that, under microaerobic conditions, the Mtr pathway could be used to drive intracellular reduction of O_2_ to water by cytochrome *bd* ^42^, which would generate a proton motive force and in turn make ATP. In these approaches, as well as others, the modularity and molecular-level specificity of the Mtr pathway allows rational design of strategies to precisely target electronic control of intracellular processes with minimal off-target effects – a long-sought goal of bioelectronics.

Our work elucidates several key points on how the MtrCAB module guides electrons out of and into *Escherichia coli* (**Figure 6**), but leaves additional points to be clarified. We demonstrate the MtrCAB conduit is needed and that electrons only reach the intracellular electron acceptor via specific oxidoreductase (Figures 2, 3). However, it is unclear how electrons transverse the periplasmic space. In these strains, MtrA is both more abundant than MtrC and is present in the periplasm, making it plausible that MtrA shuttles between MtrC and oxidoreductases. Alternatively, a native *E. coli* protein may be serving as electron carrier to a *S. oneidensis* cyt *c*. Testing these possibilities will be the subject of future work.

## Conclusions

We demonstrate here that the *mtrCAB* genetic module delivers electrons from a cathode to specific oxidoreductases so that reductions can be driven stoichiometrically in a non-native host. This finding opens new opportunities to modulate key biological processes with electrodes using a strategy that can be extended to many microorganisms.

## Materials and Methods

Additional information can be found in the Supplementary Information.

### Plasmids and strains

The strains, plasmids, primers and double stranded DNA fragments used in this study are listed in Tables S1, S2, S3 and S4, respectively. All strains were constructed using *Escherichia coli* strain C43(DE3) (Lucigen, Madison, WI). The deletion of *frdABCD, sdhABCD, nuoH, menA* and *menC* from the C43(DE3) genome was achieved using CRISPR/Cas9 (similar to Pyne et al ^43^) or λ-red recombination^44^. The pEC86 ^45^ and I5049 ^17^ plasmids carrying the *E. coli ccm* and *S. oneidensis cymAmtrCAB*, respectively, have been described previously. The pAF-*frdABCD*, pAF-*menC*, and I5105 plasmids were constructed for this work using standard molecular cloning strategies (see Supplementary Information for additional details). *E. coli* strains were grown and prepared for inoculation into bioelectrochemical reactors using standard methods (see Supplementary Information).

### Electrochemical measurements of *E. coli* strains in bioreactors

All electrochemical measurements were performed in potentiostat-controlled (VMP300, Bio-Logic LLC), three-electrode, custom-made bioelectrochemical reactors (Adams & Chittenden, Berkeley, CA) that used a cation exchange membrane (CMI-7000, Membranes International, Ringwood, NJ) to separate two 250 mL chambers. The working electrode was a 25×25 mm piece of graphite felt with a piece of Ti wire threaded vertically and the counter electrode was a piece of Ti wire. For reference we used a 3M Ag/AgCl reference electrode (CHI111, CH Instruments, Austin). The working and the reference electrode were placed in one chamber and the counter electrode was placed in the second chamber. Both chambers were filled with 140 mL M9 media and autoclaved at 121°C for 20 min. After autoclaving, filter-sterilized solutions of vitamins, minerals, amino acids, 50 mgL^-1^ kanamycin, and 10 µM IPTG were added to the working electrode chamber.

Throughout the experiment, the environmental conditions within the bioreactors were carefully controlled. The bioreactors were incubated at 30^0^C. The working electrode chamber was continuously sparged with N_2_ gas and was stirred using a magnetic stirrer rotating at ∼200 rpm. The working electrode was biassed to +0.200 V_Ag/AgCl_ and current was monitored using a potentiostat (VMP300, Bio-Logic LLC). After the baseline current stabilized (∼4 h), *E. coli* cells in fresh M9 were introduced into the bioreactor to a final cell density of 0.6 OD_600_. After 3 days of incubation (unless otherwise noted), the working electrode potential was switched to −0.560 V_Ag/AgCl_. Each experiment was replicated across three technical replicates and two biological replicates.

Spent media and cell samples were removed from the bioreactors for subsequent analysis. Spent media was collected from the working electrode chamber using a sterile needle. These samples were centrifuged at 5,000 g for 5 min to pellet any planktonic cells, and the supernatant was analyzed for the presence of small molecules with HPLC (see following section). To extract cells, the bioreactors were depolarized, gently shaken to remove the cells attached to the working electrode, and the resulting suspension was analyzed for cytochrome c content via enhanced chemiluminescence and cell density via OD_600nm_ and colony forming units (refer to SI for additional details).

### Detection of Organic Acids and Ammonia

From the supernatant samples, the concentration of various organic acids were measured by HPLC (Agilent, 1260 Infinity), using a standard analytical system (Shimadzu, Kyoto, Japan) equipped with an Organic Acid Analysis column (Bio rad, HPX-87H ion exclusion column) at 35 °C. The eluent was 5 mm sulfuric acid, used at a flow rate of 0.6 mL min^-1^ and compounds were detected by refractive index. A five-point calibration curve based on peak area was generated and used to calculate concentrations in the unknown samples. For determination of ammonia and concentrations, we employed assay kits (Sigma, AA0100) according to the manufacturer’s protocols.

## Acknowledgements

We thank Prof. Michaela TerAvest, Prof. Jeff Gralnick and Dr. Joshua Atkinson for helpful conversations and Prof. Danielle Tullman-Ercek and Kersh Thevasundaram for help with the stress-responsive promoters. Work at the Molecular Foundry was supported by the Office of Science, Office of Basic Energy Sciences, of the U.S. Department of Energy under Contract No. DE-AC02-05CH11231. This work was supported by the Office of Naval Research, Award number N000141310551.

## Author Contributions

M.B. contributed to strains construction, design of the study, conducting experiments, analysis of the data, and writing of the manuscript. S.T-S and L. S. contributed to strain construction, bioelectrochemical experiments, review and editing the manuscript. C.M.A-F. contributed to the design of the study, analysis of the data, and writing of the manuscript.

