## Supplementary Information for "Precise electronic control of redox reactions inside *Escherichia coli* using a genetic module"

**Supporting Methods:**

*Growth conditions and media composition*

All strains, unless otherwise specified, were grown in 2xYT medium at 30°C with 50 μg $mL^{-1}$ kanamycin and 30 μg $mL^{-1}$ chloramphenicol. Strains containing the pAF-*frdABCD* and pAF-*menC* plasmids were also grown with an additional 30 μg $mL^{-1}$ streptomycin. Glycerol stocks were used to inoculate 5 mL 2xYT medium, and cultures were grown overnight at 37°C with 250-rpm shaking. Then, 500 μL of overnight cultures were back-diluted into 50 mL 2xYT medium and grown in a 250 mL flask with 250-rpm shaking for 16 h at 30°C. When the cells reached an $OD_{600}$=0.5, 10 μM IPTG was added to induce production of the Mtr pathway, and the cultures were grown at 37ºC with 225 rpm shaking overnight.

The M9 media (BD) consists of 6.78 g/L disodium phosphate (anhydrous), 3 g/L $KH_2PO_4$, 0.5 g/L NaCl, 1 g/L $NH_4Cl$, and 10 mL/L each of vitamin, amino acid, and trace mineral 100x stock solutions. The 100x vitamin stock solution contained: 2 mg/L D-biotin (B7), 2 mg/L folic acid (B9), 10 mg/L pyridoxine HCl (B6), 5 mg/L thiamine HCl (B1), 5 mg/L nicotinic acid (B3), 5 mg/mL D-panthothenic acid, hexacalcium salt (B5), 0.1 mg/L cobalamin (B12), 5 mg/L p- aminobenzoic acid (PABA), and 5 mg/L α-lipoic acid. The 100x amino acid stock solution (pH 7.0) contained: 2 g/L L-glutamic acid, 2 g/L L-arginine, and 2 g/L D,L-serine. The 100x trace mineral stock solution (pH 7.0) contained: 7.85 mM $C_6H_9NO_3Na_3$, 12.17 mM $MgSO_4{\cdot}7H_2O$, 2.96 mM $MnSO_4{\cdot}H_2O$, 17.11 mM NaCl, 0.36 mM $FeSO_4{\cdot}7H_2O$, 0.68 mM $CaCl_2{\cdot}2H_2O$, 0.42 mM $CoCl_2{\cdot}6H_2O$, 0.95 mM $ZnCl_2$, 0.040 mM $CuSO_4{\cdot}5H_2O$, 0.021 mM $AlK(SO_4)_2{\cdot}12H_2O$, 0.016 mM $H_3BO_3$, 0.010 mM $Na_2MoO_4{\cdot}2H_2O$, 0.010 mM $NiCl_2{\cdot}6H_2O$, and 0.076 mM $Na_2WO_4{\cdot}2H_2O$.

The M9 media without ammonia was made from the same materials as the standard M9 medium, except $NH_4Cl$ was omitted and the final concentration of NaCl final was increased to 1.5 g/L.

*Construction of plasmids*

The plasmids used for construction of mutants are presented in Table S2, and the primers used are listed in Table S3.

To construct the pAF-*frdABCD* plasmid, we used Gibson assembly to insert the *frdABCD* operon into the pAF001 plasmid. First, we amplified the *frdABCD* operon using the primers "frdABCD+RBS (GB) fw" and "frdABCD (GB) rev" (Table S3) and using C43(DE3) genomic DNA as a template. The frdABCD+RBS (GB) fw contains both 25 bp of sequence homologous to the pAF001 plasmid and a RBS site. The pAF001 plasmid, which contains the CloDF13 ori, spectinomycin resistance cassette, and a proprionate-inducible promoter, was digested with BsaI. The digested plasmid and *frdABCD*-containing fragment were assembled together using Gibson Master Mix (New England BioLabs) as recommended by the manufacturer. The pAF-*frdABCD* plasmid was verified via sequencing.

To construct the pAF-*menC* plasmid, Gibson assembly was also used. The *menC* gene sequence was PCR amplified using primers "menC-fw" and "menC-rev" and C43(DE3) genomic DNA as the template. Both the PCR amplified gene and pAF-*frdABCD* plasmid backbone have been gel purified and then assembled via Gibson Assembly Master Mix as recommended by the manufacturer. The resulting plasmid (pAF-menC) has been PCR amplified using the primers "pAF-menC-fw" and "pAF-menC-rev" (Table S3) and verified via sequencing.

To construct the I5105 plasmid, we used a gBlock (IDT) composed of the epcD promoter (based on the sequence presented on Sergey et al 2016), RBS and the *cymA* sequence flanked by SgrAI and EcoRI digestion sites (Table S2). The gBlock was then digested by SgrAI and EcoRI. SgrAI and EcoRI were also used to digest I5049 to generate a fragment containing the plasmid backbone and *mtrCAB*. This digested fragment and the digested gBlock were ligated together and transformed into DH5ɑ competent cells (NEB) by electroporation. The resulting plasmid, I5105, containing *cymAmtrCAB* regulated by the ecpD promoter, was verified via sequencing.

*Construction of deletion strains*

**CymAMtr-*Δfrd***

The *frd* operon in C43(DE3) *E. coli* was deleted using the CRISPR-Cas9 system. We designed and obtained a gBlock (IDT) composed of a short homologous sequence of *frdA* (spacer) flanked by crispr repeats and a short homologous sequence of pMCC plasmid. The gblock was amplified using PCR and the primers "pMCC(smaI)-crisper fwd" and "pMCC(smaI)-crisper rev." The amplified gBlock was then inserted into the SmaI-digested pMCC plasmid using Gibson assembly and transformed into DH5ɑ competent cells (New England BioLabs) by electroporation. Transformants resistant to chloramphenicol were selected. The plasmid was purified from a selected clone and was sequenced verified. Next, a gBlock containing homologous DNA fragments flanking the targeted editing region was designed and obtained. This gBlock was PCR amplified with the primers "Crispr frdA-for" and "Crispr frdD-rev." Then, the pKd46-Cas9 plasmid was introduced into C43(DE3) competent cells by electroporation and transformants resistant to ampicillin were selected. The resulting C43+pKd46-Cas9 strain was grown in LB supplemented with 10 mM arabinose until $OD_{600}$=0.3-0.5 was reached. The cells were then prepared for electroporation and transformed with the pMCC plasmid and the amplified homologous DNA fragments and transformants resistant to ampicillin and chloramphenicol were selected. The knockout of the *frdABCD* operon in the selected clone has been verified via sequencing. These selected clones were grown on agar plates at $43^0$C. The last step was repeated twice, and only clones that lost resistance to chloramphenicol and ampicillin have been selected. Finally, the pEC086 and I5049 plasmids were introduced to the Δ*frdABCD* C43(DE3) background via electroporation and selection for colonies that grow on LB-agar with kanamycin and chloramphenicol.

**CymAMtr-*ΔfrdΔsdh*, CymAMtr$^s$-*ΔfrdΔsdh and* CymAMtr$^s$-*frd$^+$Δsdh***

This mutant was constructed using the lambda red mediated gene replacement, adapted from Datsenko and Wanner. The pKd46 plasmid was introduced into Mtr-*Δfrd* mutant by electroporation and transformants resistant to ampicillin were selected. The Mtr-*Δfrd*+pKd46 cells were prepared for electroporation by growing then at 30 $^0$C and adding 10mM L-arabinose when the culture reached $OD_{600}$=0.1. A DNA linear fragment containing a short 20 bp sequence homologous to the start and end of the *sdhABCD* operon and the pKd4's kanamycin resistance gene flanked by FRT sites was PCR amplified with the primers "sdh-pKd4 fw" and "sdh-pKD3 rev" (Table S3) using pKD4 as a template. The competent cells were electroporated in the presence of linear fragment and grown on a kanamycin plate at $37^0$C. Colonies containing the desired deletion of the *sdhABCD* operon were selected and verified using PCR amplification using the sdh-for and sdh-rev primers (Table S3). pCP20 was introduced into selected clones by electroporation and transformants resistant to chloramphenicol were selected. These clones were then grown again on agar plates at $43^0$C. The last step was repeated twice and a clone that lost resistance to kanamycin, chloramphenicol and ampicillin was selected. The relevant plasmids were introduced to the mutant via electroporation and selection for colonies that grow on LB-agar with antibiotics. The pEC086 and I5049 plasmids were used for the CymAMtr-*ΔfrdΔsdh* strain. The pEC086 and I5105 plasmids were used for the CymAMtr$^s$-*ΔfrdΔsdh* strain. The pEC086, I5105 and pAF-*frdABCD* plasmids were used for the CymAMtr$^s$-*frd$^+$Δsdh* strain.

**CymAMtr-*ΔnuoH and Ccm-ΔnuoH***

This mutant has been constructed using the lambda red mediated gene replacement. pKd46 plasmid was introduced into C43 (DE3) strain by electroporation and transformants resistant to ampicillin. The C43+pKd46 cells were prepared for electroporation by growing then at $30^0$C and adding 10 mM L-arabinose when the culture has reached $OD_{600}$=0.1. DNA linear fragment containing a short 20 bp sequence of the end and start of the *nuoH* gene and the pKd4's kanamycin resistance gene flanked by FRT sites was PCR amplified with The primers "nuoH-pKd3 fw" and "nuoH-pKd3 rev" (Table S3) using pKD3 as a template. The competent cells were electroporated with the linear fragment and grown on a kanamycin plate at $37^0$C. Colonies containing the desired deletion of the *nuoH* gene have been selected and verified using PCR amplification using the "nuoH-for" and "nuoH-rev" primers (Table S3). pCP20 was introduced into selected clones by electroporation and transformants resistant to chloramphenicol were selected. Cloned has been selected and grown again on agar plates at $43^0$C. The last step has been repeated twice and a clone that lost resistance to kanamycin, chloramphenicol and ampicillin has been selected. The relevant plasmids were introduced to the mutant via electroporation and selection for colonies that grow on LB-agar with antibiotics. The pEC086 and I5105 plasmids were used for the CymAMtr-*ΔnuoH* strain. The pEC086 and I5023 plasmids were used for the Ccm-*ΔnuoH* strain.

**CymAMtr-*ΔmenA***

This mutant has been constructed using the lambda red mediated gene replacement. pKd46 plasmid was introduced into C43 (DE3) strain by electroporation and transformants resistant to ampicillin. The C43+pKd46 cells were prepared for electroporation by growing then at $30^0$C and adding 10 mM L-arabinose when the culture has reached $OD_{600}$=0.1. DNA linear fragment containing a short 20 bp sequence of the end and start of the *menA* gene and the pKd4's kanamycin resistance gene flanked by FRT sites was PCR amplified with The primers "menA-pKd4-fw" and "menA-pKd4- rev" (Table S3) using pKD3 as a template. pCP20 was introduced into C43 (DE3) strain by electroporation and transformants resistant to kanamycin. Colonies containing pCP20 have been growing them at $30^0$C and screened for loss of resistance to kanamycin on plates at $37^0$C. The selected colonies were further grown on plate at $43^0$C and screened for loss of resistance to chloramphenicol. The resulting strains were tested using PCR using the primers "menA-rev" and "menA-fw"(Table S3). The pEC086 and I5049 plasmids were introduced to the mutant via electroporation and selection for colonies that grow on LB-agar with antibiotics.

**CymAMtr-*ΔmenC* and CymAMtr-$^+$*menC***

This mutant has been constructed using the lambda red mediated gene replacement. pKd46 plasmid was introduced into C43 (DE3) strain by electroporation and transformants resistant to ampicillin. The C43+pKd46 cells were prepared for electroporation by growing then at $30^0$C and adding 10 mM L-arabinose when the culture has reached $OD_{600}$=0.1. DNA linear fragment containing a short 20 bp sequence of the end and start of the *menC* gene and the pKd4's kanamycin resistance gene flanked by FRT sites was PCR amplified with The primers “menC-pKd4-fw” and “menC-pKd4- rev” (Table S3) using pKD3 as a template. pCP20 was introduced into C43 (DE3) strain by electroporation and transformants resistant to kanamycin. Colonies containing pCP20 have been growing them at $30^0$C and screened for loss of resistance to kanamycin on plates at $37^0$C. The selected colonies were further grown on plates at $42^0$C with kanamycin and ampicillin to find sensitive strains to these antibiotics. The resulting strains were tested using PCR using the primers “menC-test-rev” and “menC-test-fw”(Table S3). The relevant plasmids were introduced to the mutant via electroporation and selection for colonies that grow on LB-agar with antibiotics. The pEC086 and I5049 plasmids were used for the CymAMtr-*ΔmenC* strain. The pEC086, I5049 and pAF-menC plasmids were used for the CymAMtr-$^{+}$*menC.*

The pEC086 and I5049 plasmids were introduced to the mutant via electroporation and selection for colonies that grow on LB-agar with antibiotics.

*Analysis of cytochromes c using enhanced chemiluminescence.*

To analyze the abundance of c-type cytochromes in different *E. coli* strains, whole cell lysates were first prepared under conditions which preserve the bond between a heme *c* and cysteine residue in the protein backbone. Cells from 1.5 mL of culture were harvested by centrifugation and immediately frozen at -20°C for later analysis. Immediately before analysis, the cell pellets were thawed and resuspended in 0.1 mL Bacterial Protein Extraction Reagent (B-Per, ThermoScientific, Grand Island, NY). The cells were then lysed by addition of 6 μg $mL^{-1}$ chicken egg white lysozyme (Sigma), 1 μg $mL^{-1}$ DNAase, 3.9 mM $MgSO_4$, 0.96 mM EDTA, and 0.98 mM phenylmethylsulfonyl fluoride and incubation for 30 minutes at room temperature. The total protein concentration of the resulting whole cell lysates was determined by BCA Protein Assay Kit (ThermoScientific), and the whole cell lysates were diluted in 100 mM HEPES, pH 7.4, to yield lysates with equal total protein concentrations. Whole cell lysates were then separated by SDS-PAGE under non-reducing conditions, transferred to nitrocellulose, and the peroxidase activity of hemes *c* was detected by chemiluminescence. Whole cell lysates were diluted into NuPAGE 4x Sample Buffer (Bio-Rad) and heated at 95 °C for 5 minutes. For each strain, a total of 8 μg protein was loaded into a lane of a 4-20% Tris-HCl polyacrylamide gel (Bio-Rad) and separated by electrophoresis at 200 V for 1 hour. The gel was rinsed twice in water and then equilibrated in a cold Pierce Western Transfer buffer

(ThermoScientific) for 15 minutes. The proteins were transferred to a 0.45 μm nitrocellulose membrane (Bio-Rad, Hercules, CA) at 2.5 A 25 V for 7 minutes with Trans-Blot Turbo Transfer System (Bio-Rad). Ponceau S staining was used to confirm uniform transfer across all lanes. The nitrocellulose membrane was incubated for 5 minutes in 10 mL of Pierce Pico West Enhanced Chemiluminescence substrate (ThermoScientific), a 1:1 mixture of Pico West Peroxide Solution and Luminol Enhancer solution. The chemiluminescence signal, arising from protein-bound metals, was detected using the FluorChem E system.

*Determination of colony-forming units (CFU).*

The colony forming units (cfu) concentration for each culture was measured by preparing a serial dilution of the samples in sterile PBS. 100 µL of each relevant dilution was spread on LB-agar plates with antibiotics. The plates were grown aerobically at 37 °C and the colonies were counted the next day.

*Analysis of redox processes using cyclic voltammetry***.**

The bioreactor was prepared as previously described up until the step in which the potential was switched to -0.56 $V_{Ag/AgCl}$. At this time, cyclic voltammetry was conducted before fumarate, nitrate or nitrite addition. Cyclic voltammograms were recorded in 5 cycles from −0.8 $V_{Ag/AgCl}$ to +0.8 $V_{Ag/AgCl}$ at a scan rate of 1 mV/s. After the measurement, the electrode was switched to chronoamperometry.

*Calculation of succinate production based on the measured cathodic current*

In each experiment, the current from three polarized reactors was measured. The total charge in each reactor was integrated and converted to moles of electrons. Based on the volume of the reactor (140 mL) and the that 2 electrons are needed to reduce fumarate to succinate, these moles of electrons were converted to a change in the succinate concentration. Finally, the succinate concentration was divided by 14 days (the duration of the experiment).

$$[succinate]M \times 14\ days^{-1} = \sum_{0\ days}^{14\ days} Amp \times \frac{1\ Mole\ (electrons)}{96485\ coulomb} \times \frac{1}{0.14\ Liter} \times \frac{1\ mole\ (succinate)}{2\ Mole\ (electrons)}$$

*Calculation of succinate production based on the measured cathodic current*

In each experiment, the current from three polarized reactors was measured. The total charge in each reactor was integrated and converted to moles of electrons. To calculate the amount of ammonia produced by cathodic current, we assumed there are two possible ways that the cathodic current can be utilized for ammonia production:

1) Two moles of electrons from the cathode reduce 1 mole of nitrate to nitrite via Nrf and/or Nar complexes. Then six moles of electrons from intracellular stores, e.g. NADH, are used by Nir complex to reduce 1 mole of nitrite to 1 mole of ammonia.

$$[succinate]M \times 14\ days^{-1} = \sum_{0\ days}^{14\ days} Amp \times \frac{1\ Mole\ (electrons)}{96485\ coulomb} \times \frac{1}{0.14\ Liter} \times \frac{1\ mole\ (succinate)}{2\ Mole\ (electrons)}$$

2) The cathodic current supplies all eight moles of electrons for the production of 1 mole ammonia from nitrate.

$$[ammonia]M \times 14\ days^{-1} = \sum_{0\ days}^{14\ days} Amp \times \frac{1\ Mole\ (electrons)}{96485\ coulomb} \times \frac{1}{0.14\ Liter} \times \frac{1\ mole\ (ammonia)}{8\ Mole\ (electrons)}$$

**Supplementary Figures.**

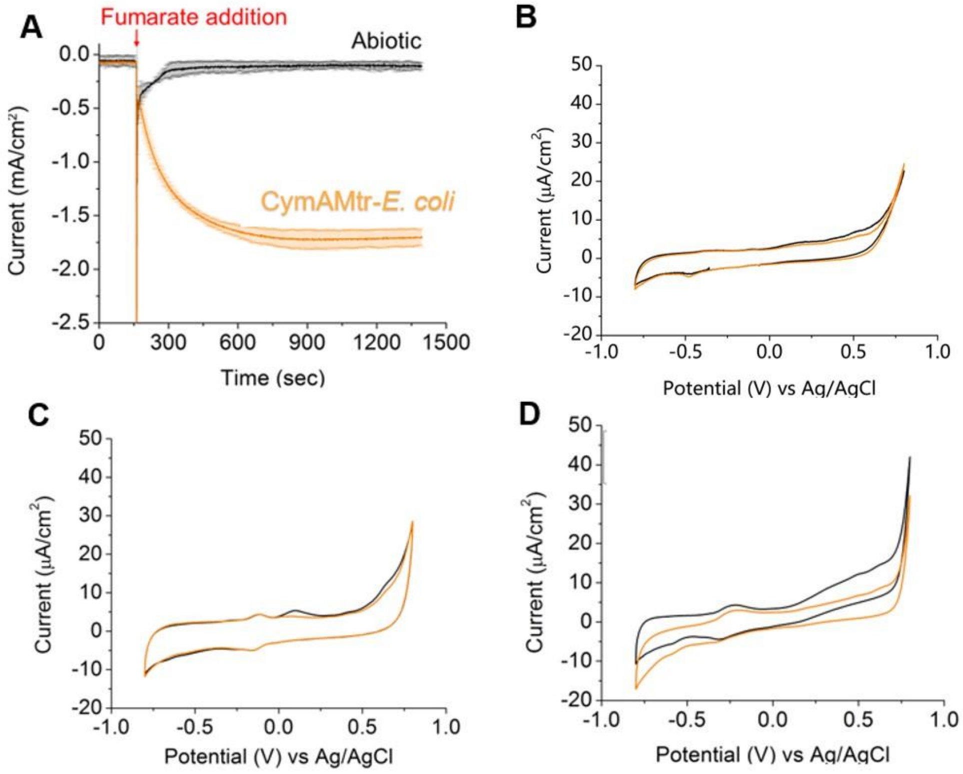


**Figure S1. Electrochemical signatures of biotic fumarate reduction.** A) Chronoamperometry upon fumarate addition to abiotic of bioelectrochemical reactors (black) and reactors containing CymAMtr-*E. coli* (orange), showing no sustained change in current in abiotic reactors. B-D) Cyclic voltammetry (CV) measurement of bioelectrochemical reactors (B) without *E. coli*, (C) the with Ccm-*E. coli* strain, and (D) with the CymAMtr-*E.coli*. Black and orange lines represent the CV before and after 50 mM fumarate addition, respectively. Only reactors containing the CymAMtr-*E. coli* show a significant catalytic wave, which is located at -350 $mV_{Ag/AgCl}$.

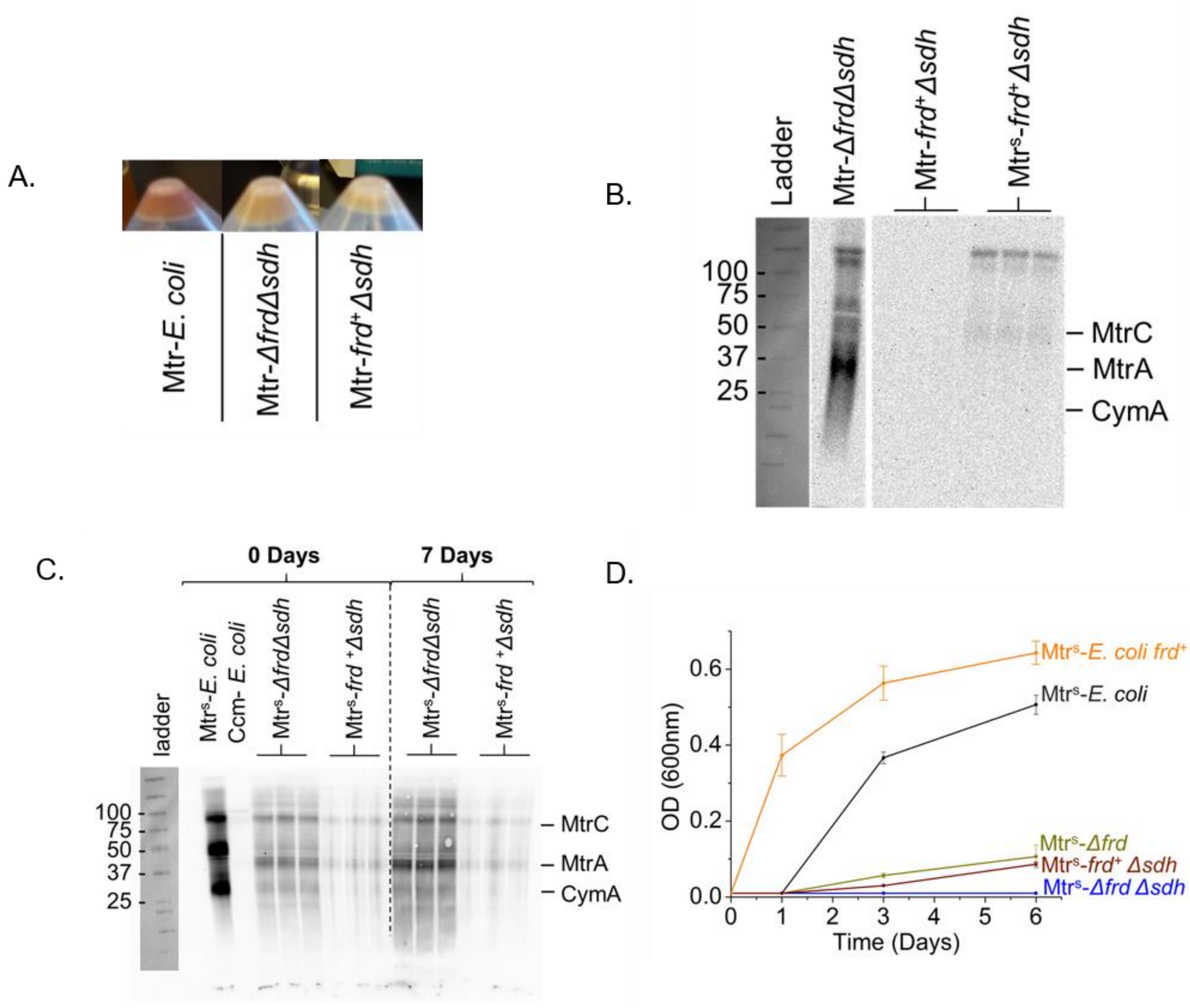


**Figure S2. Heterologous co-expression of CymAMtr and FrdABCD in *ΔfrdΔsdh* mutant.** (A) Images of *E. coli* cell pellets from the CymAMtr-*E.coli*, CymAMtr-*Δfrd Δsdh* and CymAMtr *frd*$^+$*Δsdh* after aerobic growth in 2xYT in the presence of IPTG. Expression of FrdABCD in the CymAMtr-*Δfrd Δsdh* mutant results in diminished red color of the bacteria, indicating a low abundance of matured cyts *c*. (B) Enhanced chemiluminescence (ECL) analysis of cyts *c* in the CymAMtr-*ΔfrdΔsdh*, CymAMtr-*frd*$^+$*Δsdh*, CymAMtr$^s$-*frd*$^+$*Δsdh* after aerobic growth in 2xYT in the presence of IPTG. These data indicate that introduction of a third plasmid to complement *frd* abrogates expression of the Mtr cyt *c*. However, regulating transcription of *cymAmtrCAB* by the dynamic promoter ecpD (Boyarskiy et al., 2016) restores Mtr cyt *c* expression. (C) ECL analysis of cyts *c* in the Mtr$^{s}$-*frd*$^{+}$*Δsdh* and Mtr$^{s}$-*ΔfrdΔsdh* just before inoculation into the bioelectrochemical reactors and 7 days after fumarate was added to the reactors. As a control the cyts *c* expression was examined Mtr$^{s}$-*E. coli* and Ccm-*E.coli* (the two left lanes) which were grown in the same condition as the tested strain pre inoculation. (D) Growth curves of strains grown anaerobically in minimal medium supplemented with non fermentable glycerol, as the electron donor, and fumarate as the electron acceptor.

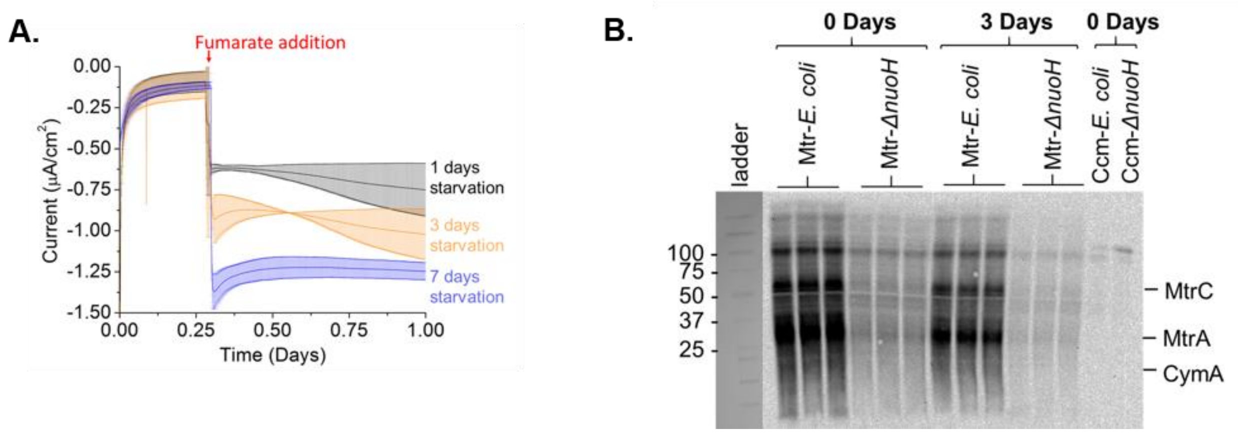


**Figure S3. The influence of starvation on current production and the effect of Complex I disruption on cyt *c* expression.** A) Chronoamperometry of CymAMtrCAB-*E. coli* upon addition of fumarate after 1 day (black), 3 days (orange), and 7 days (blue) of carbon-source deprivation, showing that increasing starvation also increases current consumption. B) ECL analysis of the CymA, MtrC, and MtrA abundance in the CymAMtr-*E. coli* and CymAMtr-*ΔnuoH* strains when inoculated into the bioelectrochemical reactor (0 days) and 3 days after after addition of fumarate (3 days). As a negative control, the expression level of Ccm-*E. Coli* and Ccm-*ΔnuoH* strain is shown. Those strains were grown in the same condition as the tested strain pre-inoculation.

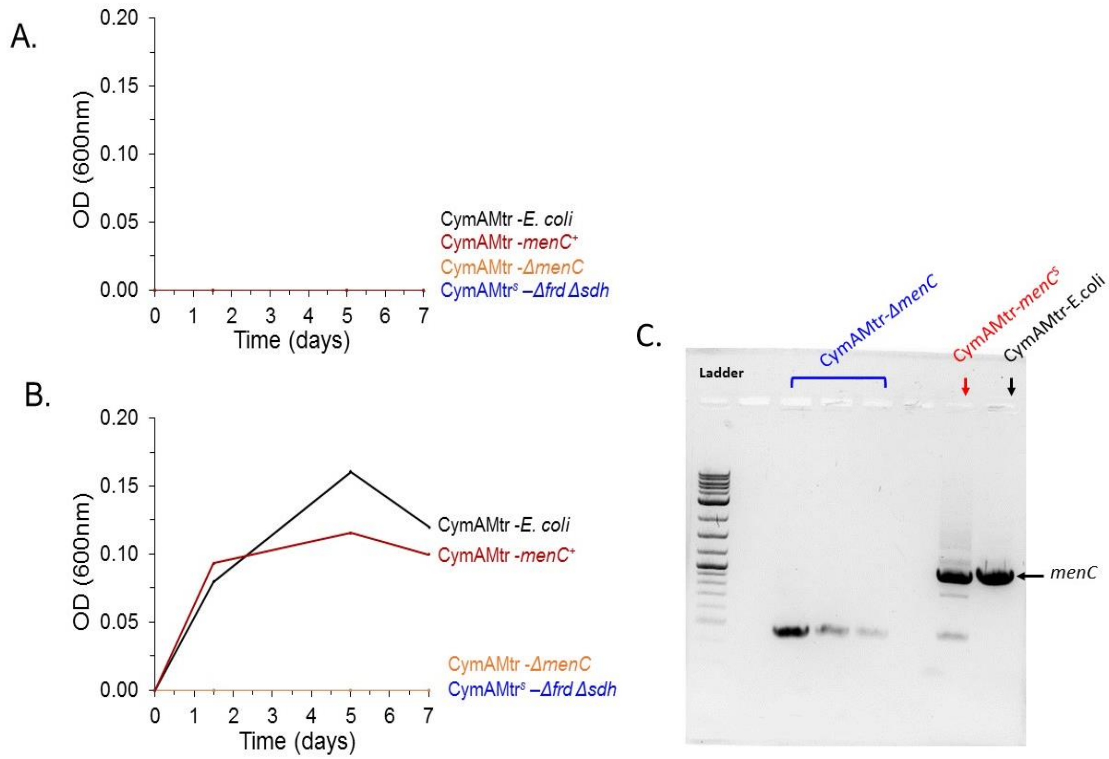


**Figure S4. MenC is essential for fumarate respiration under anaerobic condition in CymAMtr-*E. coli* strain .** A-B**)** Growth curves of strains grown in an anaerobic in minimal medium containing glycerol as the electron donor. No electron acceptor is provided (A) or Fumarate is provided as an electron acceptor (B). C) Agarose gel electrophoresis of the PCR products obtained from the genomic DNA from CymAMtr-E.coli, CymAMtr-*ΔmenC* and CymAMtr-*menC*$^{S}$ strains.

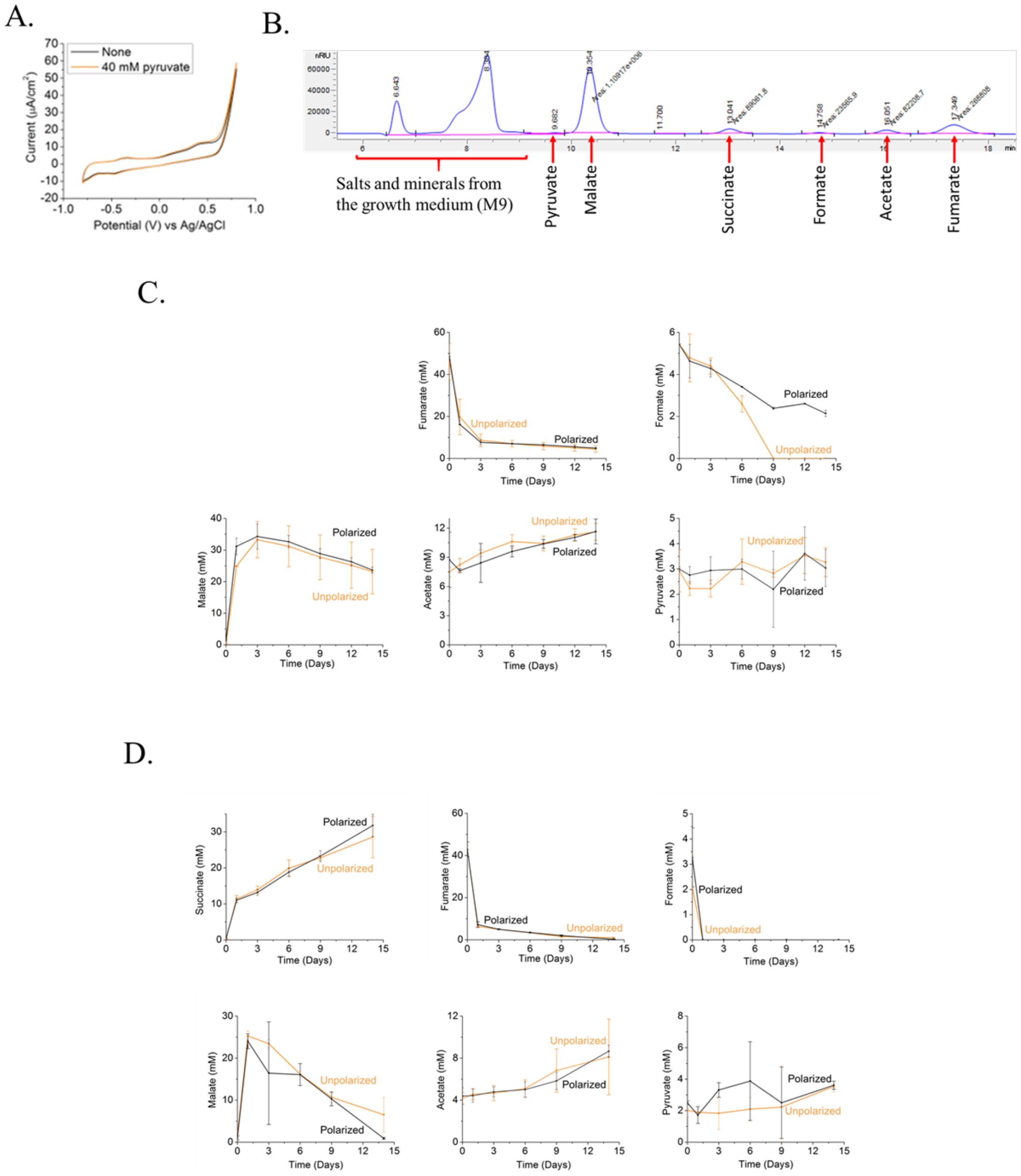


**Figure S5.** (A) Cyclic voltammetry measurement showing no sustained change in current between abiotic reactors that don't contain or contain 40mM pyruvate (B) Representative HPLC chromatogram of a sample from a the supernatant of a bioreactor after 14 days of Mte-E. coli incubation. The chromatogram displays the peaks of the following analytes: pyruvate, malate, succinate, formate, acetate and fumarate. (C) A plot representing the supernatant acetate, pyruvate malate and fumarate concentration in Bias (Black) vs unbiased (orange) bioreactors containing the

Mtr-*ΔnuoH* mutant. Measurement started upon 50mM Fumarate addition and were taken over a period of 14 days values of 0.0 on plots indicating that the concentration was below the detection limit in that sample. All values are means of triplicate bioreactors, and error bars represent standard error. (D) A plot representing the supernatant succinate, formate, acetate, pyruvate malate and fumarate concentration in Bias(Black) vs unbiased (orange) bioreactors containing the Mtr-*E. coli* strain. Measurement started upon 50mM Fumarate addition and were taken over a period of 14 days values of 0.0 on plots indicating that the concentration was below the detection limit in that sample. All values are means of triplicate bioreactors, and error bars represent standard error.

**Supplementary Tables.**

**Table S1. Strains used in this study.**

| **Nickname** | **Description** | **Strain #** | **Genotype** | **Source** |
|---|---|---|---|---|
| Ccm-*E.coli* | *E. coli* expressing *ccmABCDEFGH* and carrying an empty Kan vector | MFe408 | C43(DE3) +pEC86, pSBIET2 | Goldbeck et al. 2013. |
| Mtr-*E.coli* | *E. coli* expressing IPTG-inducible *mtrCAB* | MFe409 | C43(DE3) +pEC86, I5023 | Goldbeck et al. 2013. |
| CymAMtr-*E.coli* | *E. coli* expressing IPTG-inducible *cymAmtrCAB* | MFe444 | C43(DE3) +pEC86, I5049 | Jensen et al. 2016. |
| CymAMtr- *Δfrd* | *E. coli* expressing IPTG-inducible *cymAmtrCAB* in the *ΔfrdABCD* background | MFe1085 | C43(DE3) *ΔfrdABCD* +pEC86, I5049 | This study |
| CymAMtr-*ΔfrdΔsdh* | *E. coli* expressing IPTG-inducible *cymAmtrCAB* in the *ΔfrdABCD ΔsdhABCD* background | MFe1086 | C43(DE3) *ΔfrdABCD ΔsdhABCD* +pEC86, I5049 | This study |
| CymAMtr$^{S}$-*ΔfrdΔsdh* | *E. coli* expressing *cymAmtrCAB* under control of the *epcD* promoter in the *ΔfrdABCD ΔsdhABCD* background | MFe854 | C43(DE3) *ΔfrdABCD ΔsdhABCD* +pEC86, I5105 | This study |
| CymAMtr$^{S}$-*frd*$^{+}$*Δsdh* | *E. coli* expressing *cymAmtrCAB* under control of the *epcD* promoter and *frdABCD* in the *ΔfrdABCD ΔsdhABCD* background | MFe1089 | C43(DE3) *ΔfrdABCD ΔsdhABCD* + I5105+pEC86+pAF-frdABCD | This study |
| CymAMtr-*ΔmenA* | *E. coli* expressing IPTG-inducible *cymAmtrCAB* in the *ΔmenA* background | MFe1083 | C43(DE3) *ΔmenA* +pEC86, I5049 | This study |
| CymAMtr-*ΔmenC* | *E. coli* expressing IPTG-inducible *cymAmtrCAB* in the *ΔmenC* background | MFe1084 | C43(DE3) *ΔmenC* +pEC86, I5049 | This study |

| | | | | |
|---|---|---|---|---|
| CymAMtr-*menC*$^+$ | *E. coli* expressing IPTG-inducible *cymAmtrCAB* and *menC* in the *ΔmenC* background | MFe1128 | C43(DE3) *ΔmenC* +pEC86, I5049, pAF-menC | This study |
| Ccm-*ΔnuoH* | *E. coli* constitutively expressing *ccmABCDEFGH* in the *ΔnuoH* background | MFe1088 | C43(DE3) *ΔnuoH* +pEC86, pSB1ET2 | This study |
| CymAMtr-*ΔnuoH* | *E. coli* expressing IPTG-inducible *cymAmtrCAB* in the *ΔnuoH* background | MFe1087 | C43(DE3) *ΔnuoH* +pEC86+I5049 | This study |

**Table S2. Plasmids used in this study.**

| **Plasmid** | **Description** | **Source** |
|---|---|---|
| pEC86 | Constitutive expression of *ccmABCDEFGH* | Goldbeck et al. 2013 (Goldbeck et al., 2013) |
| I5049 | IPTG-inducible expression of *cymAmtrCAB* | Jensen et al 2016 (Jensen et al., 2016) |
| I5023 | IPTG-inducible expression of *mtrCAB* | Jensen et al 2010 (Jensen et al., 2010) |
| I5105 | expression of *cymAmtrCAB* under control of Stress-responsive promoter *epcD* promoter | This study |
| pSB1ET2 | *Empty backbone of I5049* | Jensen et al 2010 (Jensen et al., 2010) |
| pAF-frdABCD | *Expression of frdABCD* | This study |
| pCP20 | *Expression* the *Flp* recombinase | Cherepanov et al 1995 (Cherepanov & Wackernagel, 1995) |
| pAF-MenC | *Expression of menC* | This study |

**Table S3.** Primers Used in This Study, Related to the Experimental Procedures

| Primer name | Sequence 5’ -> 3’ | Reference |
|---|---|---|
| sdh-pKD3 fw | 'cgatagcgtccattctccatcgcgtttccggtgtgatcacctttgttgcaggtgtaggctggagctgcttc' | This study |
| *sdh-pKD3 rev* | 'gatggcgcgcgtcgggttcagccccttcggacatacactgacgcagttcaatgggaattagccatggtcc' | This study |
| *sdh-rev* | ‘atgcttacttcgccgtggat’ | This study |
| *sdh-fw* | ‘tgaacagcctatactgccgc’ | This study |
| nuoH-pKd3 fw | 'ctgaccatcctcaaagcggtggtgatcctgctggtggttgtcacctgcgggtgtaggctggagctgcttc' | This study |
| nuoH-pKd3 rev | 'tattgcgcctgccagagaatgacagccgccgttaccagcaagttgatcagatgggaattagccatggtcc' | This study |
| nuoH- fw | tccggtgctggctggcgcgcatcttgagga | This study |
| nuoH- rev | gcaggccgatcatccagata | This study |
| *frdABCD+RBS – fw (GB)* | 'aataacaagttgataacaagctagccaaaaacaacattctaacta' | This study |

| | | |
|---|---|---|
| frdABCD – rev (GB) | 'ttcgttttatttgatgcctggttattttacattggcgatgcgtta' | This study |
| pMCC(SmaI)-crispr.fwd | ‘atacgcaaaccgcctctccccatggatcctatttcttaataactaaaaatatgg' | This study |
| pMCC(SmaI)-crispr.rev | 'gaagaacccttccagtgccccactgagacttgttgagttgaattcatg' | This study |
| Crisper frdA-for | 'gtgcaaacctttcaagccga' | This study |
| Crisper frdD-rev | 'aggatagcagccagaccgta' | This study |
| menA-pKd4-fw | ‘atgactgaacaacaaattagccgaactcaggcgtggctggaaagtttacgaccgtgt’ | This study |
| menA-pKd4-rev | ‘ttatgctgcccactggcttaggaatatccctaaaacaaacagcaggttagtatggga’ | This study |
| menA-fw | ‘tatttgtcagttatgcygcccactggcttaggaa’ | This study |
| menA-rev | ‘acacactgttctggagcgtttaatggaa’ | This study |
| menC-pKd4-fw | ‘tctccccgcacttgccgccagtgacgccacggccagtcagagaagatcatgtgtagg’ | This study |
| menC-pKd4-rev | ‘tcaaccagaaacgtcagcctgacttcagcaaattcaaacggaatccgtaaatgggaa’ | This study |
| menC-test-fw | ‘agcatttgctccaatggctgt’ | This study |

| | | |
|---|---|---|
| menC-test-rev | ‘cttgtgaacaccgtggtacc’ | This study |
| menC-fw | ‘attctaactacggttactctatgcgtagcgcgcaggtata’ | This study |
| menC-rev | ‘tattttacattggcgatgcgtcataacaaccgctccagtgc’ | This study |
| pAF-menC-fw | ‘ccaccagtgccggaatagtt’ | This study |
| pAF-menC-rev | ‘atcgtccggatcgccattac’ | This study |

**Table S4.** gBlocks Used in This Study, Related to the Experimental Procedures

| **gblock name** | **Sequence 5’ -> 3’** | **Reference** |
|---|---|---|
| p*epcD*-RBS-*cymA* | ‘ccgatcttccccatcggtgatgtcggcgatataggcgccagcaa ccgcacctgtggcgccggtgatgccggccacgatgcgtccggc gtagaggatcgagatcgatctcgatctaaagccctggatgcaacg acaaaatctgcttatgattttaaagcgtcttacgttcgtgccgtcgca gaccaaacagcaactgctggttatgtaaaaactaaaataatttctatt ttatattattccctgttttaattaactctatcagggatggtttgtttaactt taagaaggagatatacatattttggagatagagtaatgaactggcg tgcactatttaaacccagcgcgaaatattccatcctagcgctactgg ttgttggtatcgtgattggtgttgtgggctattttgcaactcagcaga ctttacatgcgacaagtacagatgcgttctgtatgtcttgccatagca atcattccttgaagaatgaagtgctggcatctgcccacggtggcgg caaagccggggttactgttcagtgtcaagactgtcacttaccccat ggccctgttgattatttaattaagaaaatcatcgtatctaaagatttata tggtttcttaactattgatggctttaacactcaagcttggttagacgaa | This study |

| | | |
|---|---|---|
| | aaccgcaaagagcaagccgacaaagcattggcttacttccgtggt aacgactcagcaaactgtcaacactgccatactcgcatttatgaaa accagccagaaaccatgaagccaatggctgtgagaatgcacacc aacaacttcaagaaagatcctgaaacgagaaagacctgtgtggat tgccacaaaggtgtcgctcacccctatccaaaaggataaggtttaa cgctgcaag’ | |
| *frdA*-crisper | 'catggatcctatttcttaataactaaaaatatggtataatactcttaat aaatgcagtaatacaggggcttttcaagactgaagtctagctgaga caaatagtgcgattacgaaattttttagacaaaaatagtctacgagg ttttagagctatgctgttttgaatggtcccaaaacatgacccaactgg aactgtgggggtgcccagttttagagctatgctgttttgaatggtcc caaaacttcagcacatgaattcaactcaacaagtctcagtg' | This study |
